# Modelling complex population structure using *F*-statistics and Principal Component Analysis

**DOI:** 10.1101/2021.07.13.452141

**Authors:** Benjamin M Peter

**Affiliations:** MPI Evolutionary Anthropology

## Abstract

Human genetic diversity is shaped by our complex history. Data-driven methods such as Principal Component Analysis (PCA) are an important population genetic tool to understand this method. Here, I contrast PCA with a set of statistics motivated by trees (*F*-statistics). Here, I show that these two methods are closely related, and I derive explicit connections between the two approaches. I show that *F*-statistics have a simple geometrical interpretation in the context of PCA, and that orthogonal projections are the key concept to establish this link. I illustrate my results on two examples, one of local, and one of global human diversity. In both examples, I find that just using the first few PCs provides good population structure is sparse, and only a few components contribute to most statistics. Based on these results, I develop novel visualizations that allow for investigating specific hypotheses, checking the assumptions of more sophisticated models. My results extend *F*-statistics to non-discrete populations, moving towards more complete and less biased descriptions of human genetic variation.

## 1 Introduction

As in most species, the genetic diversity of human populations has been influenced by our history and environment over the last several hundred thousand years (e.g Cavalli-Sforza et al., 1994; Marciniak and Perry, 2017; Reich, 2018; Nielsen et al., 2017; Witt et al., 2022). In turn, an important goal of population genetics is to use observed patterns of variation to investigate and reconstruct the demographic and evolutionary history of our species (Schraiber and Akey, 2015; Orlando et al., 2021).

The complicated genetic structure observed in present-day human populations (The 1000 Genomes Project Consortium, 2015; Mallick et al., 2016) is caused by the interplay of demographic and evolutionary processes with both discrete and continuous components (Pritchard et al., 2000; Rosenberg et al., 2002; Serre and Pääbo, 2004; Rosenberg et al., 2005; Bradburd et al., 2018; Reich, 2018; Peter et al., 2020; Gopalan et al., 2022). In particular, populations are expected to slowly differentiate if they are isolated from each other (Wahlund, 1928; Cavalli-Sforza and Piazza, 1975). In humans, this may be caused because continental-scale geographic distances limit migration, causing a pattern known as isolation-by-distance (Slatkin, 1985). However, these patterns are usually not uniform, but shaped by geography, particularly barriers to migration such as mountain ranges, oceans or deserts (Cavalli-Sforza et al., 1994; Barbujani and Sokal, 1990; Rosenberg et al., 2005; Bradburd et al., 2013; Peter et al., 2020). In addition, major historical population movements such as the out-of-Africa, Austronesian or Bantu expansions lead to more gradual patterns of genetic diversity over space (Cavalli-Sforza et al., 1994; Ramachandran et al., 2005; Novembre et al., 2008; Stoneking, 2016; Racimo et al., 2020). Local migration between neighboring populations will reduce differentiation, and long-distance migrations (Alves et al., 2016), and secondary contact between diverged populations, such as Neandertals and modern humans (Green et al., 2010) may lead to locally increased diversity (Gopalan et al., 2022).

Particularly for large and heterogeneous data sets, disentangling all these processes is challenging, and we cannot expect to devise a single model catching both broad strokes and minute details of human history. A commonly used analysis paradigm is thus to integrate tools based on different sets of assumptions. each emphasizing particular aspects of the data.

A typical analysis starts with data-driven, exploratory methods that summarize data making minimal assumptions (e.g. Schraiber and Akey, 2015). Examples are population trees (Cavalli-Sforza and Edwards, 1967; Felsenstein, 1973; Cavalli-Sforza and Piazza, 1975), Principal Component Analysis (PCA, Cavalli-Sforza et al., 1994; Patterson et al., 2006)) structure-like models (Pritchard et al., 2000; Alexander et al., 2009) or multidimensional scaling (MDS Lessa, 1990)). However, these methods are not designed to answer specific research questions, and are limited in their ability to estimate biologically meaningful parameters. For this purpose, methods based on explicit demographic models are often used that aim to fit a specified or estimated model of divergence, migration and genetic drift to the data (Gutenkunst et al., 2009; Excoffier et al., 2013; Kamm et al., 2015). The drawback of these methods is that, to make inference mathematically feasible, we need to introduce strong modeling assumptions such as that populations are discrete, randomly mating, or at equilibrium. While in most cases these assumptions are violated to some extent and cannot be verified, we hope that the resulting model fits provide sufficiently accurate answers to specific research questions.

However, when the number of populations exceeds a few dozen, even codifying reasonable population models can be prohibitively difficult. One approach is to pick a small set of “representative” samples, and restrict modeling to this subset (e.g. Gravel et al., 2011; Harney et al., 2021). However, this has the drawback that a large proportion of the available data remains unused. An increasingly popular alternative approach, particularly in the analysis of human ancient DNA, is therefore to build up complex models from smaller building blocks based on the relationship between two, three or four populations.

The framework is based on a set of parameters called *F*-statistics *sensu* Patterson (Reich et al., 2009; Patterson et al., 2012; Peter, 2016). Formal definitions will be given in the Theory section; but an informal motivation starts with the null model that populations are related as a tree, in which each *F*-statistic measures the length of a particular set of branches. (Figure 1; Semple and Steel, 2003; Peter, 2016).

**Figure 1:**
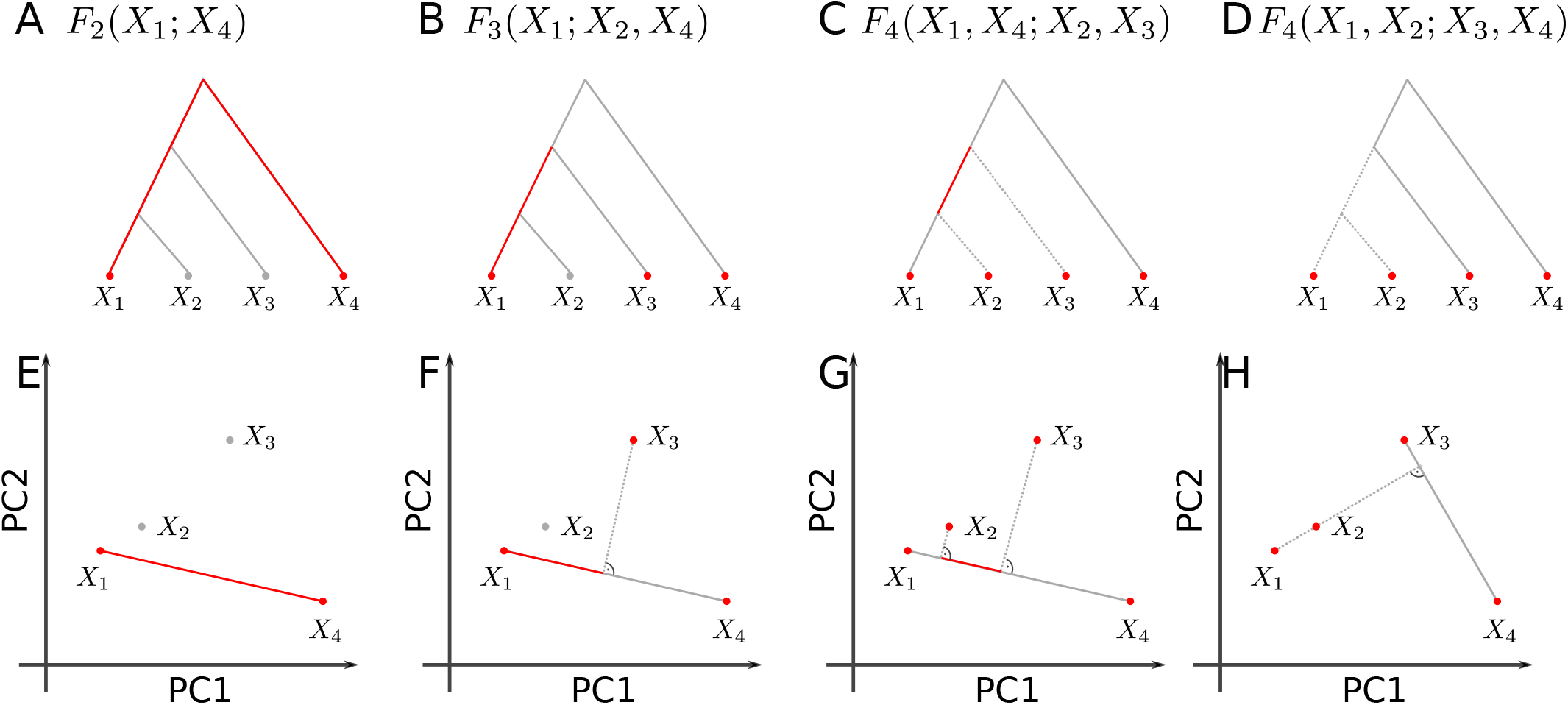
Representation of *F*-statistics on trees and 2D-PCA-plots. The schematics show four populations and their representation using a tree (top row) or a 2D-PCA plot (bottom row). *A*: *F*_2_ represents the (squared) Euclidean distance between two tree leafs, and in PC-space. B: *F*_3_(*X*_1_; *X*_3_, *X*_4_) corresponds to the external branch from *X*_1_ to the internal node joining the populations, and is proportional to the orthogonal projection of *X*_1_ – *X*_3_ onto *X*_1_-*X*_4_. C: *F*_4_(*X*_1_, *X*_4_; *X*_2_, *X*_3_) corresponds to the internal branch in the tree, or the orthogonal projection of *X*_2_ – *X*_3_ on *X*_1_ – *X*_4_. D: *F*_4_(*X*_1_, *X*_2_; *X*_3_, *X*_4_) The two paths from *X*_1_ to *X*_2_ and *X*_3_ and *X*_4_ are non-overlapping in the tree, which corresponds to orthogonal vectors in PCA-space.

In most applications, *F*-statistics are estimated from data, and then used as tests of treeness. In particular, under the assumption of a tree, *F*_3_ is restricted to be non-negative, and many *F*_4_-statistics will be zero (Semple and Steel, 2003; Patterson et al., 2012), and data that violates these constraints is incompatible with a tree-like relationship between populations. The canonical alternative model is an admixture graph (or phylogenetic network) (Patterson et al., 2012; Huson et al., 2010), which is a tree which allows for additional edges reflecting gene flow (Figure 2A). However, admixture graphs are not the only plausible alternative models, and expected *F*-statistics can be calculated for a wide range of population genetic demographic models (Peter, 2016).

**Figure 2:**
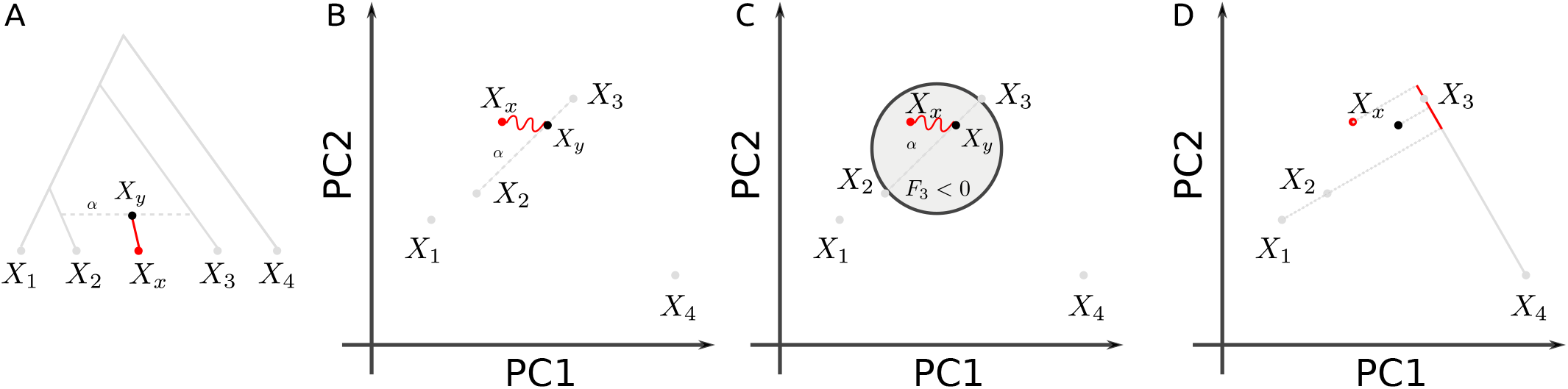
Admixture representation on 2D-PCA-plot. The schematics show four populations and their representation using an admixture graph (A) or a 2D-PCA plot. A: Admixture graph, with population *X_y_* originating as an admixture of *X*_2_ and *X*_3_, with *X*_2_ contributing proportion α. Subsequent drift (red branch) will change allele frequency to sampled admixture population *X*_x_. B: PCA representation of the scenario in A. *X*_y_ originates on the segment connecting *X*_2_ and *X*_3_, and subsequent drift may move it in a random direction. C: *F*_3_(*X_x_*; *X*_2_, *X*_3_) and negative region (light gray circle). *F*_4_(*X*_1_, *X_x_*; *X*_3_, *X*_4_) will no longer be zero (compare to Figure 1D).

### F-statistics and PCA

The practical issue addressed in this study is how *F*-statistics can be reconciled with PCA, one of the most widely used data-driven modeling techniques. One way PCA can be motivated is as generating a low-dimensional representation of the data, with each dimension (called principal component, PC) retaining a maximum of the variance present in the data. To understand population structure, the use of PCA has been pioneered by Cavalli-Sforza et al., 1964, who used allele-frequency data at a population level to visualize genetic diversity (Cavalli-Sforza et al., 1994). Currently, PCA is most commonly performed on individual-level genotype data (e.g. Patterson et al., 2006; Novembre et al., 2008), making use of the hundreds of thousands of loci available in most genome-scale data sets. The PCA-decomposition has been studied for a number of explicit population genetic models including trees (Cavalli-Sforza and Piazza, 1975), spatially continuous structure (Novembre and Stephens, 2008), the coalescent (McVean, 2009) and discrete population models (François and Gain, 2021). Here, in order to link PCA to *F*-statistics, I interpret both of them geometrically in *allele frequency space,* i.e. as functions of a high-dimensional Euclidean space. For *F*-statistics, this interpretation was recently developed by Oteo-Garcia and Oteo, 2021, and for PCA it follows naturally from the interpretation of approximating a high-dimensional space with a low-dimensional one.

In the next section, I will formally derive the connection between *F*-statistics and PCA, and show how *F*-statistics can be interpreted geometrically, with a particular emphasis on two-dimensional PCA plots. In the Results section, I will then discuss how some of the most common applications of *F*-statistics manifest themselves on a PCA, and illustrate them on two example data sets, before ending with a discussion.

## 2 Theory

In this section, I will introduce the mathematics and notations for *F*-statistics and PCA. A comprehensive treatise on PCA is given by e.g. Jolliffe, 2013, a useful primer on the mathematics is Pachter, 2014, and a helpful guide to interpretation is Cavalli-Sforza et al., 1994. Readers unfamiliar with *F*-statistics may find Patterson et al., 2012, Peter, 2016 or Oteo-Garcia and Oteo, 2021 helpful.

### 2.1 Formal Definition of *F*-statistics

Let us assume we have a set of populations for which we have SNP allele frequency data from *S* loci. Let *x_il_* denote the frequency at the *l*-th SNP in the *i*-th population; and let *X_i_* = (*x*_*i*1_, *x*_*i*2_,… *x_iS_*) be a vector collecting all allele frequencies for population *i*. As *X_i_* will be the only data summary considered here for population *i*, I make no distinction between the population and the allele frequency vector used to represent it.

The three *F*-statistics are defined as

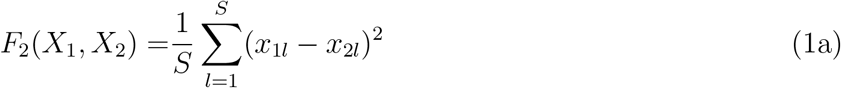

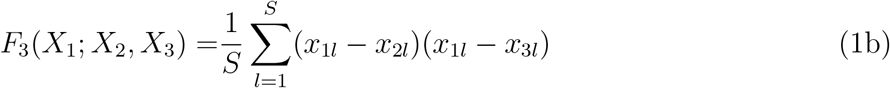

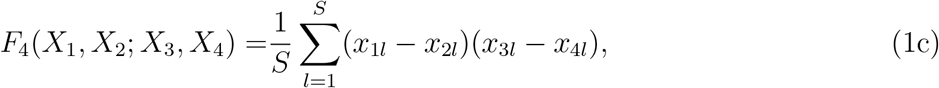

The normalization by the number of SNPs *S* is assumed to be the same for all calculations and is thus omitted subsequently. Both *F*_3_ and *F*_4_ can be written as sums of *F*_2_-statistics:

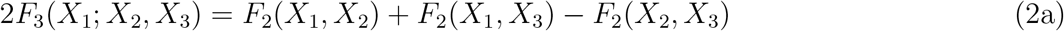

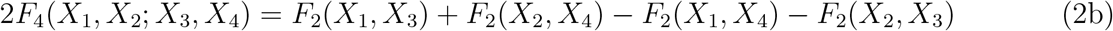

*F*-statistics have been primarily motivated in the context of trees and admixture graphs (Patterson et al., 2012). In a tree, the squared Euclidean distance *F*_2_(*X*_4_, *X*_2_) measures the length of the path between populations *X*_4_ and *X*_2_ (Figure 1A); *F*_3_ represents the length of an external branch (Figure 1B) and *F*_4_ the length of an internal branch, respectively (Figure 1C). Crucially, for branches that do not exist in the tree (as in Figure 1D), *F*_4_ will be zero. The length of each branch can be thought of in units of genetic drift, and is non-negative (Patterson et al., 2012).

Thinking of *F*-statistics as branch lengths is useful for a number of applications, including building multi-population models (Patterson et al., 2012; Lipson et al., 2013), estimating admixture proportions (Petr et al., 2019; Harney et al., 2021) and finding the population most closely related to an unknown sample (“Outgroup”-*F*_3_-statistic).

Most commonly however, *F*_3_ and *F*_4_ are used as tests of treeness (Patterson et al., 2012): Negative *F*_3_-values correspond to a branch with negative genetic drift, which is not allowed under the null assumption of a tree-like population relationship. Similarly if four populations are related as a tree, then at least one of the *F*_4_ statistics between the populations will be zero (Buneman, 1974; Patterson et al., 2012).

The most widely considered alternative model is an admixture graph (Patterson et al., 2012), an example is given in Figure 2A. Here, the (typically unobserved) population *X_y_* is generated by a mixture of individuals from the ancestors of *X*_2_ and *X*_3_. Over time, genetic drift will change *X_y_* to *X_x_*, which is the admixed population we observe. This will result in *F*_4_-statistics that are non-zero, and, in some cases, in negative *F*_3_-statistics (exact conditions can be found in Peter, 2016).

#### 2.1.1 Geometric interpretation of *F*-statistics

An implicit assumption in the development of *F*-statistics is that population lineages are mostly discrete, and that gene flow is rare. Recently, Oteo-Garcia and Oteo, 2021 re-derived *F*-statistics in a geometric framework, showing that these assumptions are not necessary. Specifically, they interpret the populations *X_i_* as points or vectors in the *S*-dimensional *allele frequency space* ℝ^*S*^. In this case, the *F*-statistics can be thought of as inner (or dot) products, and they showed that all properties and tests related to treeness can be derived in this larger space. In particular the *F*-statistics can be written as

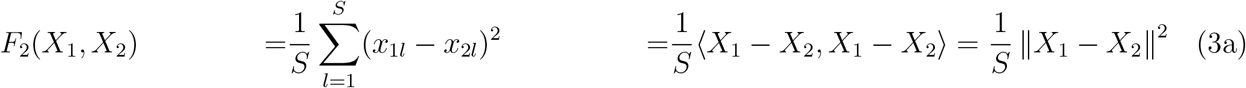

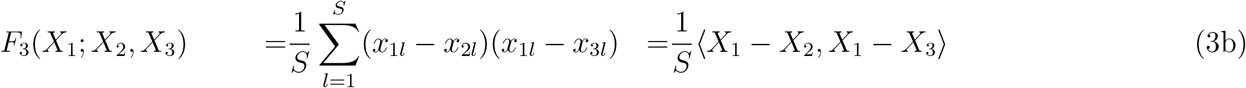

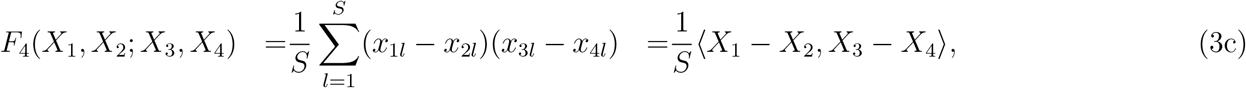

where ||·|| denotes the Euclidean norm and 〈·, ·〉 denotes the dot product. Some elementary properties of the dot product between vectors *a, b, c* that I will use later are

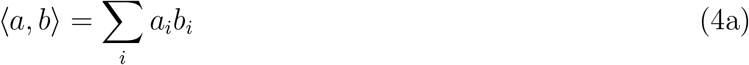

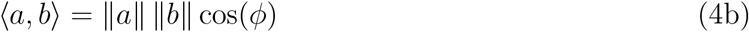

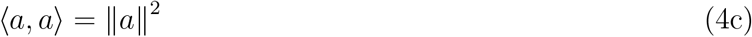

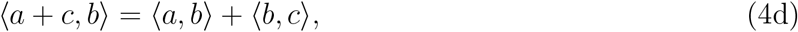

where *ϕ* is the angle between *a* and *b*. The inner product is closely related to the vector projection

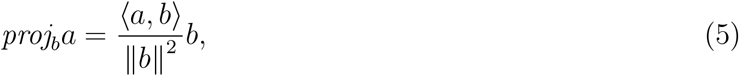

which is a vector colinear to *b* whose length measures how much vector a points in the direction of *b*. Thinking of *F*-statistics as projections also holds on trees: In e.g. a *F*_4_(*X*_1_, *X*_4_; *X*_2_, *X*_3_)-statistic (Figure 1C), the internal branch is precisely the intersection of the paths from *X*_1_ to *X*_4_ and from *X*_2_ and *X*_3_. On trees, all disjoint paths are independent (i.e. orthogonal) from each other, and thus the external branches vanish under the projection.

The drawback of the geometric approach of Oteo-Garcia and Oteo, 2021 is that we have to deal with an very high-dimensional space, as the number of SNPs is frequently in the millions. However, it has been commonly observed that population structure is quite low-dimensional, and that the first few PCs provide a good approximation of the covariance structure in the data (Patterson et al., 2006). Therefore, we may hope that PCA could yield a reasonable approximation of the allele frequency space, and that *F*-statistics as measures of population structure may likewise be well-approximated by the first few PCs.

### 2.2 Formal Definition of PCA

PCA is a common way of summarizing genetic data, and so a large number of variations of PCA exist, e.g. in how SNPs are standardized, how missing data is treated, what algorithm is used, or whether we use individuals or populations as units of analysis. The version of PCA I use here is set up such that the similarities to *F*-statistics are maximized, and does *not* reflect how PCA is most commonly applied to genome-scale human genetic variation data sets. In particular, I assume that a PCA is performed on unscaled, estimated population allele frequencies. In contrast, many applications of PCA are based on individual-level sample allele frequency, scaled by the estimated standard deviation of each SNP (Patterson et al., 2006). The differences this causes will be addressed in the discussion.

Let us again assume we have allele frequency data as above, but let us now assume we aggregate the allele frequency vectors *X_i_* of many populations in a matrix **X** whose entry *x_il_* reflects the allele frequency of the *i*-th population at the *l*-th genotype. If we have *S* SNPs and n populations, **X** will have dimension *n* × *S*. Since the allele frequencies are between zero and one, we can interpret each population *X_i_* of **X** as a point in ℝ^*S*^.

PCA allows us to approximate the points in the high-dimensional allele frequency space by a *K*-dimensional subspace of the data. If all PCs are considered, *K* = *n* – 1, in which case the data is simply rotated. However, the historical processes that generated genetic variation often result in *low-rank* data (Engelhardt and Stephens, 2010), so that *K* ≪ *n* explains a substantial portion of the variation; for visualization *K* = 2 is frequently used.

There are several algorithms that are used to perform PCAs, the most common one is based on singular value decomposition (e.g. Jolliffe, 2013). In this approach, we first mean-center **X**, obtaining a centered matrix **Y**

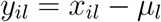

where *μ_l_* is the mean allele frequency at the *l*-th locus.

PCA can then be written as

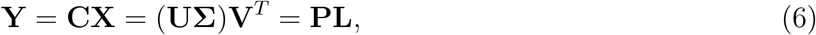

where 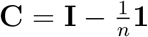 is a centering matrix that subtracts row means, with **I, 1** the identity matrix and a matrix of ones, respectively. For any matrix **Y**, we can perform a singular value decomposition **Y** = **UΣV**^*T*^ which, in the context of PCA, is interpreted as follows: The matrix of principal components **P** = **UΣ** has size *n* × *n* and contains information about population structure. The SNP loadings **L** = **V**^*T*^ form an orthonormal basis of size *n* × *S*, its rows give the contribution of each SNP to each PC. It is often used to look for outliers, which might be indicative of selection (e.g Duforet-Frebourg et al., 2016). Alternatively, the PCs can also be obtained from an eigendecomposition of the covariance matrix **YY**^*T*^. This can be motivated from (6):

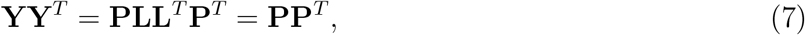

since **LL**^*T*^ = **I**.

### 2.3 Connection between PCA and *F*-statsitics

#### 2.3.1 Principal components from *F*-statistics

PCA, as defined above, and *F*-statistics are closely related. In fact, the principal components can be directly calculated from *F*-statistics using multidimensional scaling, which, for squared Euclidean (*F*_2_)-distances, leads to an identical decomposition to PCA (Gower, 1966). Suppose we calculate the pairwise *F*_2_(*X_i_,X_j_*) between all *n* populations, and collect them in a matrix **F**_2_. We can obtain the principal components from this matrix by double-centering it, so that its row and column means are zero, and perform an eigendecomposition of the resulting matrix:

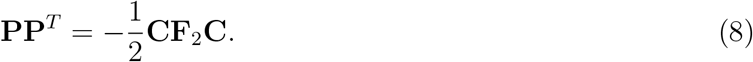

#### 2.3.2 *F*-statistics in PCA-space

By performing a PCA, we rotate our data to reveal the axes of highest variation. However, the dot product is invariant under rotation, and *F*-statistics can be thought of as dot products (Oteo-Garcia and Oteo, 2021). What this means is that we are free to calculate *F*_2_ either on the uncentered data ***X***, the centered data **Y** or any other orthogonal basis such as the principal components **P**. Formally,

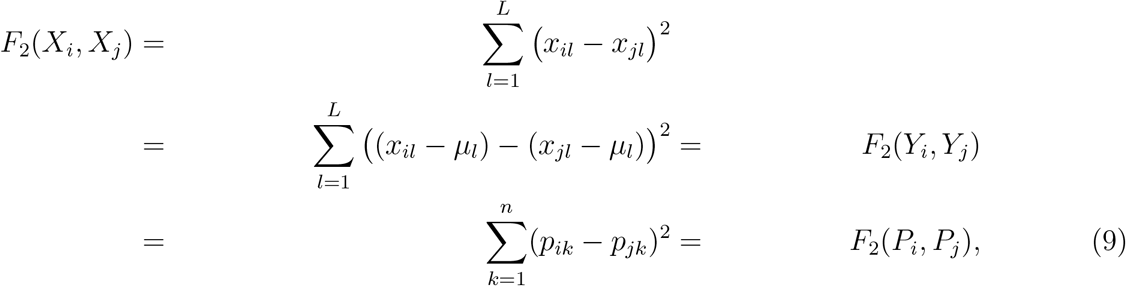

A derivation of this change-of-basis is given in Appendix A, Equation A1. As *F*_3_ and *F*_4_ can be written as sums of *F*_2_-terms (Eqs. 2a, 2b), analogous relations apply.

In most applications, we do not use all PCs, but instead truncate to the first *K* PCs, which explain most of the between-population genetic variation. Thus,

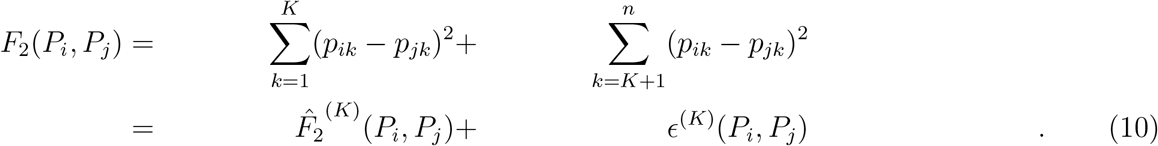

In this notation, 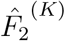 is the approximation of *F*_2_ with only the first K PCs considered, and *ϵ*^(*K*)^ is the corresponding approximation error. I will omit the superscript of 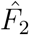 when the exact number of PCs is not relevant. If we sum up the squared approximation errors over all pairs of populations in our sample, we obtain

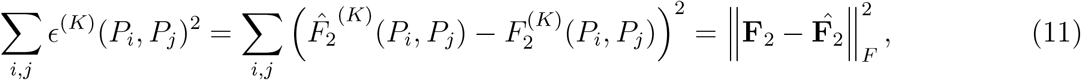

where the Frobenius-norm 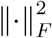 of a matrix is defined as the square root of the sum-of-squares of all its elements. This is precisely the function that is minimized in MDS (Jolliffe, 2013). In that sense, 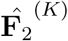 is the optimal low-rank approximation of **F**_2_ for any *K* in that it minimizes the sum of approximation errors of all *F*_2_-statistics.

#### 2.3.3 *F*-statistics and samples projected onto PCA

One of the easiest ways of dealing with missing data in PCA is to calculate the principal components (equation 6) only on a subset of the data with no missingness, and then to *project* the lower quality samples with high missingness onto this PCA. The simplest way to do this is to note that

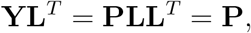

and so a new (centered) population Y_*new*_ can be projected onto an existing PCA simply by postmultiplying it with **L**^*T*^:

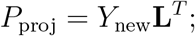

the *k*-th entry of *P*_proj_ gives the coordinates of the new sample on the *k*-th PC. However, it is likely that *Y*_new_ lies outside the variation of the original samples. In this case, there is a projection error

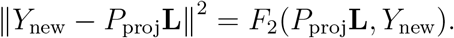

If we project with missing data, a similar projection can be used where we remove the rows from *Y*_new_ and **L** where data in *Y*_new_ is missing, and add a scaling factor (Patterson et al., 2006).

Thus, if we compare the *F*-statistic of a projected sample, we have

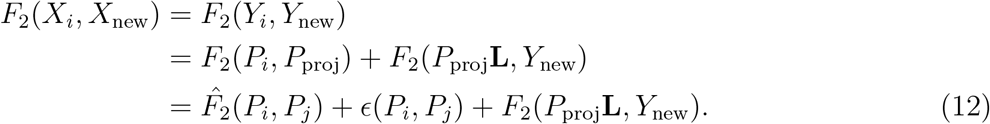

The second row follows because the projection error and projection are orthogonal to each other. The main implication of equation 12 is that both truncation and projection introduce some error, and that 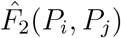 will be a good approximation to *F*_2_(*P_i_, P_j_*) only if both errors are small.

## 3 Material & Methods

The theory outlined in the previous section suggests that *F*-statistics have a geometric interpretation in PCA-space, which can be approximated on PCA plots. In the next section I explore this connection in detail, and illustrate it on two sample data sets that I briefly introduce here. Both are based on the analyses by Lazaridis et al., 2014. The data is from the Reich lab compendium data set (v44.3), downloaded from https://reich.hms.harvard.edu/allen-ancient-dna-resource-aadr-downloadable-genoo using data on the “Human Origins”-SNP set (597,573 SNPs). SNPs with missing data in any population are excluded. The code used to write this paper, create all figures and analyses is available on https://github.com/BenjaminPeter/fstats_pca.

### “World” data set

This data set is a subset of the “World Foci” data set of Lazaridis et al., 2014, where I removed samples which are not permitted for free reuse. These populations span the globe and roughly represents global human genetic variation (638 individuals from 33 population) As adjacent sampling locations are often thousands of kilometers apart, I speculate that gene flow between these populations may not be particularly common; and their structure may therefore be well-approximated by an admixture graph. A file with all individuals used and their assigned population is given in **Supplementary File 1.**

### Western Eurasian data set

This data set of 1,119 individuals from 62 populations contains present-day individuals from the Eastern Mediterranean, Caucasus and Europe. Lazaridis et al., 2014 used this data set as a basis of comparison for ancient genetic analyses of Western Eurasian individuals, and PCAs based on similar sets of samples have been used in many other ancient DNA studies (e.g. Lazaridis et al., 2016; Haak et al., 2015). Genetic differentiation in this region is low and closely mirrors geography (Novembre et al., 2008). I thus speculate that gene flow between these populations is common (Ralph and Coop, 2013), and a discrete model such as a tree or an admixture graph might be a rather poor reflection of this data. A file with all individuals used and their assigned population is given in **Supplementary File 2.**

### Computing *F*-statistics and PCA

All computations are performed in `R`. I use `admixtools` 2.0.0 (https://github.com/uqrmaie1/admixtools) to compute *F*-statistics. To obtain a PC-decomposition, I first calculate all pairwise *F*_2_-statistics, and then use equation 8 and the eigen function to obtain the PCs. The right-hand side matrix of equation 8 is supposed to have non-negative eigenvalues (i.e. –**CF**_2_**C** is positive-semidefinite). However as *F*_2_-statistics are estimates, some eigenvalues might be slightly negative, which would lead to imaginary PCs. I avoid this by setting all negative eigenvalues to zero.

## 4 Results

The transformation from the previous section allows us to consider the geometry of *F*-statistics in PCA-space. The relationships we will discuss formally only hold if we use all PCs. However, the appeal of PCA is that frequently, only a very small number *K* ≪ *n* of PCS contain most information that is relevant for population structure, in which case the geometric interpretations become very simple. Thus, throughout the schematic figures, I assume that two PCs are sufficient to characterize population structure. In the data applications I evaluate how deviations of this assumption may manifest themselves in PCA plots.

### 4.1 *F*_2_ in PC-space

The *F*_2_-statistic is an estimate of the squared allele-frequency distance between two populations. On a tree (Figure 1A) this corresponds to the branch between two populations (Figure 1E). In allele-frequency space, it corresponds to the squared Euclidean distance, and thus reflects the intuition that closely related populations will fall close to each other on a PCA-plot, and have low pairwise *F*_2_-statistics. However, since *F*_2_ can be written as a sum of squared (non-negative) terms for each PC (eq. 9), the distance on a PCA-plot will always be an underestimate of the full *F*_2_-distance. Thus, PCA might project two populations with high *F*_2_-distance very close to each other, which would indicate that these particular PCs are not suitable to understand and visualize the relationship between these particular populations, and likely more PCs need to be investigated to understand how these populations are related to each other. In converse, populations that are distant on a PCA-plot are guaranteed to also have a large *F*_2_-distance.

### 4.2 When are admixture-*F*_3_ statistics negative?

Consider again the admixture scenario in Figure 2A, where population *X_y_* is the result of a mixture of *X*_2_ and *X*_3_, and subsequent drift changes the allele frequencies of the admixed population from *X_y_* to *X_x_*. How is such a scenario displayed on a PCA? Since the allele frequencies of *X_y_* are a linear combination of *X*_2_ and *X*_3_, it will lie on the line segment connecting these two populations (Figure 2B), at a location predicted by the admixture proportions. Subsequent drift will change the allele frequency of *X_x_*, and so in general it might fall on a different point on a PCA-plot. An exception occurs when *X_x_* (and no other populations related to *X_x_*) are not part of the construction of the PCA, so that *X_x_* – *X_y_* is orthogonal to all PCs, i.e.

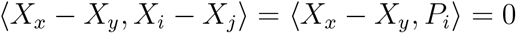

for all populations *i, j* ≤ *n*. In this case, *X_x_* and *X_y_* project to the same point, and the location on the PCA can directly be used to predict the admixture proportions (McVean, 2009; Brisbin et al., 2012; Oteo-Garcia and Oteo, 2021). However, if either *X_x_*, is included in the construction of the PCA, or if some gene flow occurred between *X_x_* and any of the populations used to construct the PCA, *X_x_* and *X_y_* may project on different spots (Figure 2B).

Thus, a natural question to ask is given two source populations *X*_2_, *X*_3_, can we use PCA to predict which populations might have negative *F*_3_-statistics? This condition can be written as

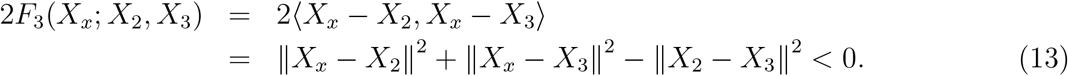

By the Pythagorean theorem, *F*_3_ = 0 if and only if *X*_2_, *X*_3_ and *X_x_* form a right-angled triangle. The associated region where *F*_3_ = 0 is a *n*-sphere (or a circle in two dimensions) with diameter 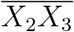 (The overline denotes a line segment). *F*_3_ is negative when the triangle is obtuse, i.e. *X_x_* could be considered admixed if it lies inside the *n*-ball with diameter 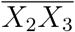 (Figure 1B, Equation A2).

#### *F*_3_ on a 2D PCA-plot

If we project this *n*-ball on a two-dimensional plot, 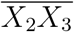 will usually not align with the PCs; thus the ball may be somewhat larger than it appears on the plot. This geometry is perhaps easiest visualized on a globe. If we look at the globe from a view point parallel to the equator, both the north and south poles are visible at the very edge of the circle. But if we look at it from above the north pole, the north- and south-poles will be at the very same point.

Thus if 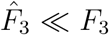, the “true” circle will be bigger than what would be predicted from a 2D-plot, and populations that appear inside the circle on a PCA-plot may, in fact, have positive *F*_3_-statistics. This is because they are outside the *n*-ball in higher dimensions. The converse interpretation is more strict: if a population lies outside the circle on *any* 2D-projection, *F*_3_ is guaranteed to be bigger than 0 (see Equation A4 in the Appendix).

#### Example

As an example, I visualize the admixture statistic *F*_3_(*X*; Sardinian, Finnish), on the first two PCs of the Western Eurasian data set (Figure 3A). In this case, the projected *n*-ball (light gray) and circle based on two dimensions (dark gray) have similar sizes. However, several populations that appear inside the circles (e.g. Basque, Canary Islanders) have, in fact, positive *F*_3_-values, so they lie outside the *n*-ball. This reveals that the first two PCs do not capture all the genetic variation relevant for European population structure. Consequently, approximating *F*_3_ by the first two or even ten PCs (Figure 3B) only gives a coarse approximation of *F*_3_, and from Figure 3C we see that many higher PCs contribute to *F*_3_ statistics.

**Figure 3:**
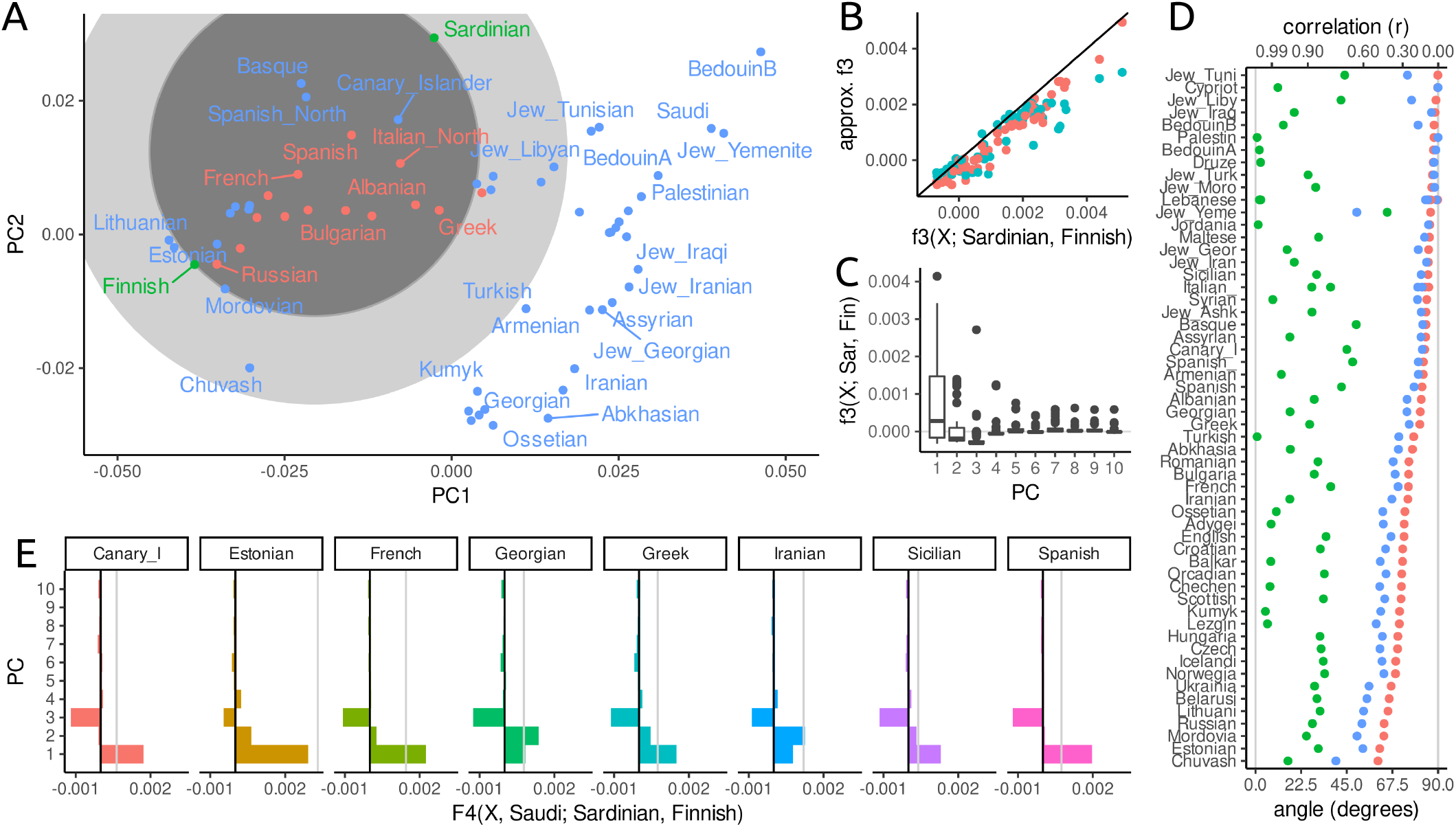
PCA and *F*-statistics for the Western Eurasian data set. A: PCA-biplot; the light grey circle denotes the region for which *F*_3_(*X*; Sardinian, Finnish) may be negative, the dark circle is based on just the first two PCs. Populations for which *F*_3_ is negative are colored in red. B: *F*_3_ approximated with two (blue) and ten (red) PCs versus the full spectrum. C: Boxplot of contributions of PCs 1-10 to each *F*_3_-statistic. D: Projection angle and correlation interpretation of *F*_4_(*X*, Saudi; Sardinian, Finnish) based on two PCs (green), three PCs (blue) or full data (red). E: Contribution of the first ten PCs to select *F*_4_-statistics, with the first three PCs containing the majority of contributions.

However, many populations, particularly from Western Asia and the Caucasus, on the right-hand side of the plot, fall outside the circle. This allows us to immediately conclude that their *F*_3_-statistics must be positive; and we should not consider them as a mixture between Sardinians and Fins.

### 4.3 Outgroup-*F*_3_-statistics as projections

A common application of *F*_3_-statistics is, given an unknown sample *X_U_*, to find the most closely related population among a reference panel (*X_i_*) (Raghavan et al., 2014). This is done using an *outgroup*-*F*_3_-statistic *F*_3_(*X_O_*; *X*_U_,*X_i_*), where *X_O_* is an outgroup. The reason an outgroup is introduced is to account for differences in sample times and additional drift in the reference populations (Figure 4A). The outgroup-F_3_-statistic *F*_3_(*X_O_*; *X_U_*, *X*_3_) represents the branch length from *X_O_* to the common node between the three samples in the statistic, and the closer this node is to *X_U_*, the longer the branch and hence the larger the *F*_3_-statistic.

**Figure 4:**
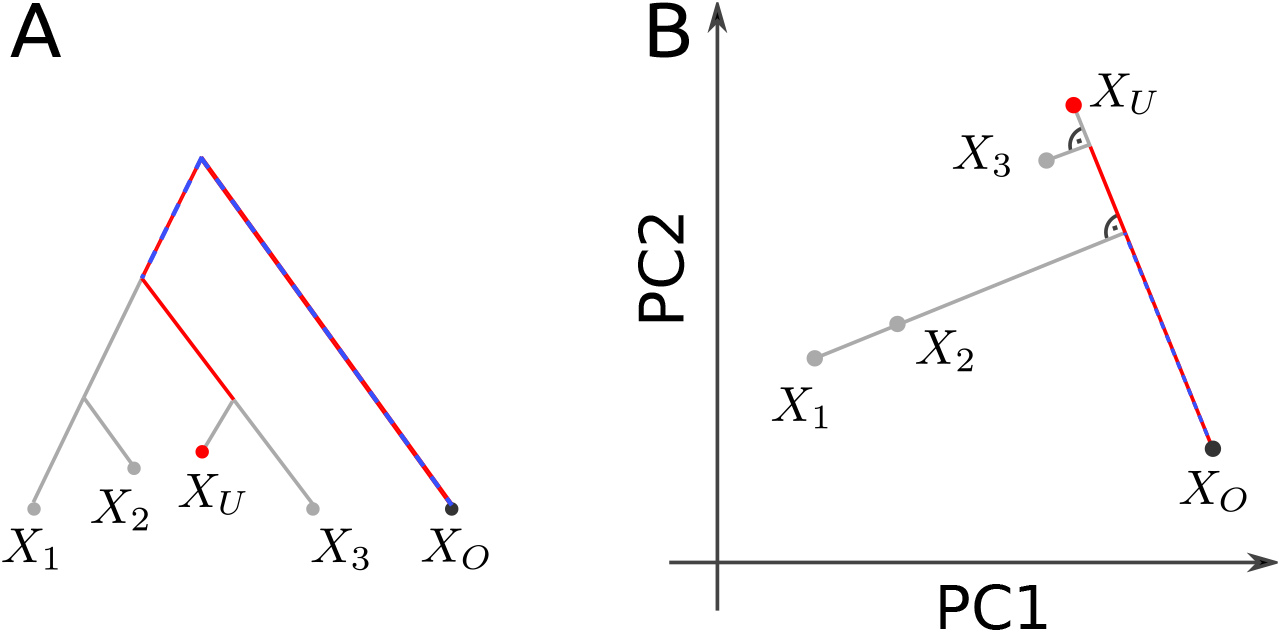
Outgroup-*F*_3_-statistics. Interpretation of outgroup *F*_3_-statistic on a tree (A) and PCA-plot (B). The red segment represents *F*_3_(*X_O_*; *X_U_*, *X*_3_) and the dashed blue segment reflects *F*_3_(*X_O_*; *X_y_*, *X_i_*) and *F*_3_(*X_O_*; *X_U_*, *X*_2_), which have the same value.

To make sense of outgroup-*F*_3_-statistics in the PCA context, I use the association of *F*_3_-statistics to projections (Equation 5): On a PCA-plot, we can visualize this *F*_3_-statistic as the projection of the vector *X_i_* – *X_O_* onto *X_U_* – *X_O_*:

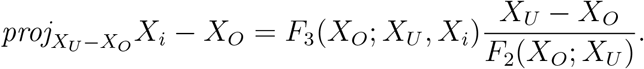

Of the right-hand-side terms, only the *F*_3_ term depends on the *X_i_*. The fraction can be thought of as a normalizing constant, so the *F*_3_-statistic is proportional to the length of the projected vector, and they can thus be interpreted similarly. This means, that the outgroup-*F*_3_-statistic is largest for whichever *X_i_* projects furthest along the axis from the outgroup to the unknown population; in Figure 4 this is *X*_3_.

#### Example

In Figure 5A, I use the World data set to visualize the outgroup-*F*_3_-statistic *F*_3_(Mbuti; Sardinian, *X* i.e. a statistic that aims to find the population most closely related to Sardinian (a Mediterranean Island), assuming the Mbuti are an outgroup to all populations in the data set. On a PCA, we can interpret this *F*_3_ statistic as the projection of the line segment from Mbuti to population *X_i_* onto the line through Mbuti and Sardinians (black line). For each population, the projection is indicated with a grey line. In the full allele frequency space, this line is always orthogonal to the segment Mbuti-Sardinian, but on the plot (i.e. the subspace spanned by the first two PCs), this is not necessarily the case. The coloring is based on the *F*_3_-statistic calculated from all the data, with brighter i values indicating higher *F*_3_-statistics. In this case, the first two PCs approximate the *F*_3_-statistic very well: Particularly the samples from East Asia, Siberia and the Americas (cluster in the top left of the plot) project very close to orthogonally, suggesting that most of the genetic variation relevant for this analysis is captured by these first two PCs. We can quantify this and find that the first two PCs slightly underestimate the absolute value of *F*_3_ (Figure 3C), but keep the relative ordering. I also find that many PCs, e.g. PCs 3-5, 7 and 10 have almost zero contribution to all *F*_3_-statistics (Figure 3D), and PCs 6, 8 and 9 having a similar non-zero contribution for almost all statistics, likely because these PCs explain within-African variation.

**Figure 5:**
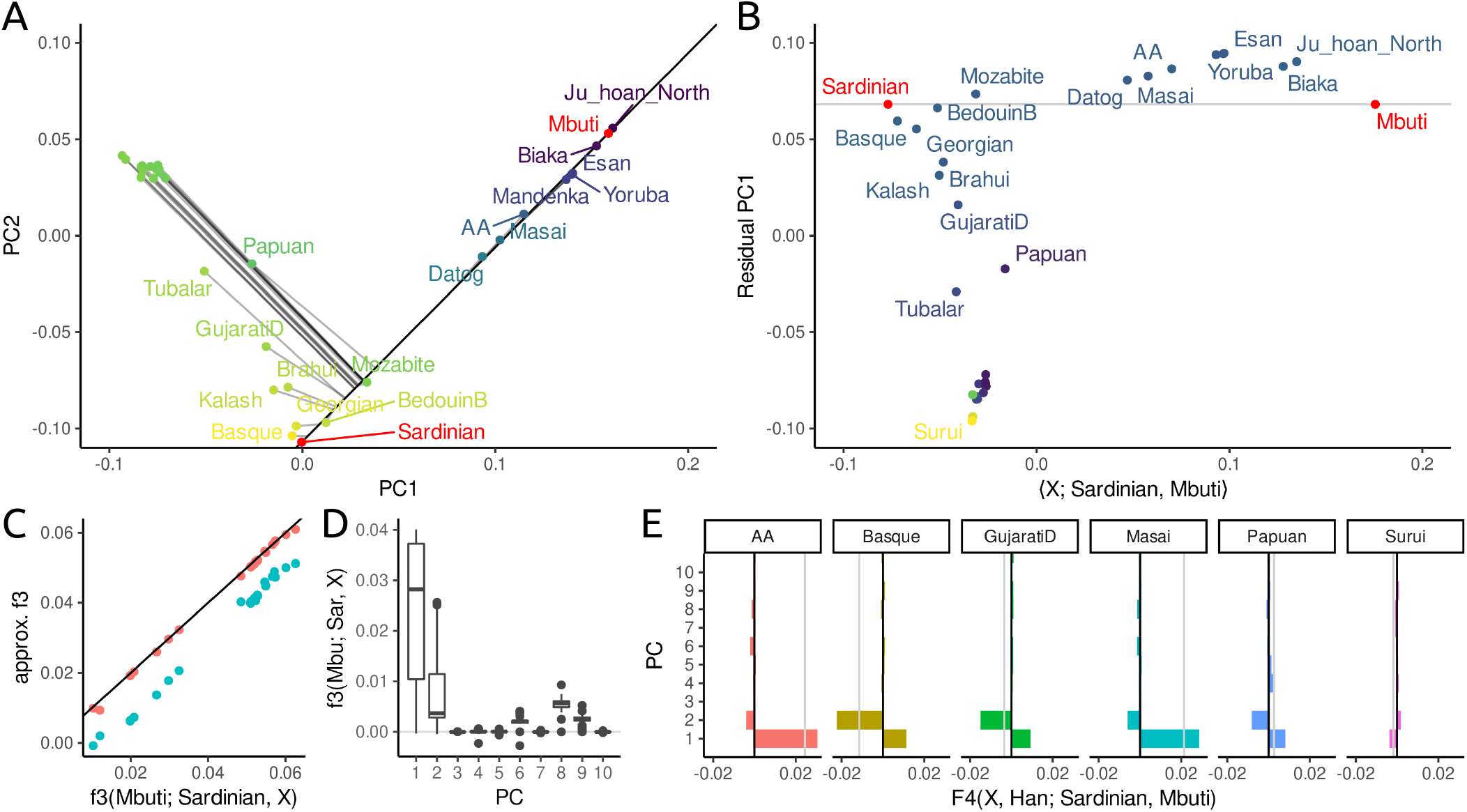
PCA and *F*-statistics for the World data set. A: Visualization of Outgroup-*F*_3_-statistic *F*_3_(Mbuti; Sardinian, *X*) on a PCA-biplot. The color of points correspond to the value of the *F*_3_ statistic, with brighter yellows indicating higher values, i.e. higher similarity to Sardinians. The *F*_3_-projection axis is given by a black line, the projection of populations onto this axis by thin gray lines. In the full allele frequency space, these projection are orthogonal to the axis. B: Projection along the axis Sardinian-Mbuti (*X*-axis), and PCA on residual of this projection (PC1 on *Y*-axis, PC2 as coloring). C: Approximation of *F*_3_(Mbuti; Sardinian, *X*) using the first two (blue) and first ten (red) PCs, respectively. D: Contributions of first ten PCs to all statistics of the form *F*_3_(Mbuti; Sardinian, *X*). E: Contributions of the first ten PCs to select *F*_4_-statistics.

### 4.4 *F*_4_-statistics as angles

One interpretation of *F*_4_ on PCA plots is similar to that of *F*_3_; as a projection of one vector onto another, with the difference that now all four points may be distinct. *F*_4_-statistics that correspond to a branch in a tree (as in Figure 1C), can be interpreted as being proportional to the length of a projected segment on a PCA plot (Figure 1G), again with the caveat that we need to scale it by a constant. If the *F*_4_-statistic corresponds to a branch that does not exist in the tree, i.e it is a test statistic (Figure 1D), then, from the tree-interpretation, we expect *F*_4_(*X*_1_, *X*_2_; *X*_3_, *X*_4_) = 0 which implies that the vectors *X*_1_ – *X*_2_ and *X*_3_ – *X*_4_ are orthogonal to each other, i.e. that *X*_1_ and *X*_2_ map to the same point on the projection axis 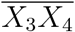 (Figure 1H). In the case of an admixture graph, this is no longer the case: Both population *X_y_* and *X_x_* in Figure 2D do *not* map to the same point as *X*_1_ or *X*_2_ do, implying that statistics of the form *F*_4_(*X*_1_, *X_x_*; *X*_3_, *X*_4_) ≠ 0.

Since *F*_4_ is a covariance, its magnitude lacks an interpretation. Therefore, commonly correlation coefficients are used, as there, zero means independence and one means maximum correlation. For *F*_4_, we can write

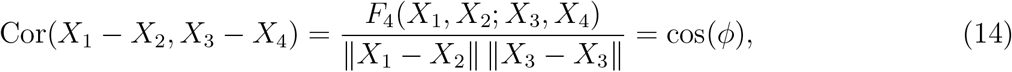

where *ϕ* is the angle between *X*_1_ – *X*_2_ and *X*_3_ – *X*_4_. Thus, independent drift events lead to cos(*ϕ*) = 0, so that the angle is 90 degrees, whereas an angle close to zero means cos(*ϕ*) ≈ 1, which means most of the genetic drift on this branch is shared.

#### Example

To illustrate the angle interpretation I return to the Western Eurasian data. The PCA-biplot shows two roughly parallel clines (Figure 3A), a European gradient (from Sardinian to Finnish and Chuvash), and a Asian cline from Arab populations (top right) to the Caucasus (bottom right). This is quantified in Figure 3D, where I plot the angle corresponding to *F*_4_(*X*, Saudi; Sardinian, Finnish). For most Asian populations, using two PCs (green points) gives an angle close to zero, corresponding to a correlation coefficient between the two clines of *r* > 0.9. Just adding a third PC (blue), however, shows that the clines are not, in fact, parallel, and the correlation for most populations is low. The finding that three PCs are necessary to explain this data can also be seen from the spectrum of these *F*_4_-statistics (Figure 3E), which have high contributions from the first three PCs.

### 4.5 Other projections

So far, I used eq. 9 to interpret *F*-statistics on a PCA-plot, but the argument holds for *any* orthonormal projection of the allele frequency space. This is useful in particular for estimates of admixture proportions, which are often done as projections into a low-dimensional reference space defined by *F*-statistics (Patterson et al., 2012; Petr et al., 2019; Harney et al., 2021; Oteo-Garcia and Oteo, 2021).

For example, a common way to estimate admixture proportion is the *F*_4_-ratio:

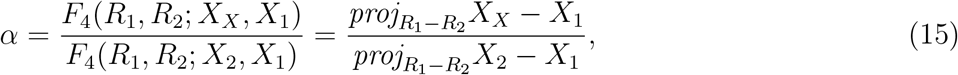

which can be interpreted as projecting *X_X_* – *X*_1_ and *X*_2_ — *X*_1_ onto *R*_1_ – *R*_2_ and the ratio of the lengths gives the proportion of *X_x_* contributed by *X*_1_ (Oteo-Garcia and Oteo, 2021).

The admixture graph motivating this statistic is visualized in Figure 6A, and the PCA-like interpretation in Figure 6B. In both panels, the solid gray lines is the projection axis, and the dotted lines give the residual, i.e. the branches or genetic variation that is ignored by the projection.

**Figure 6:**
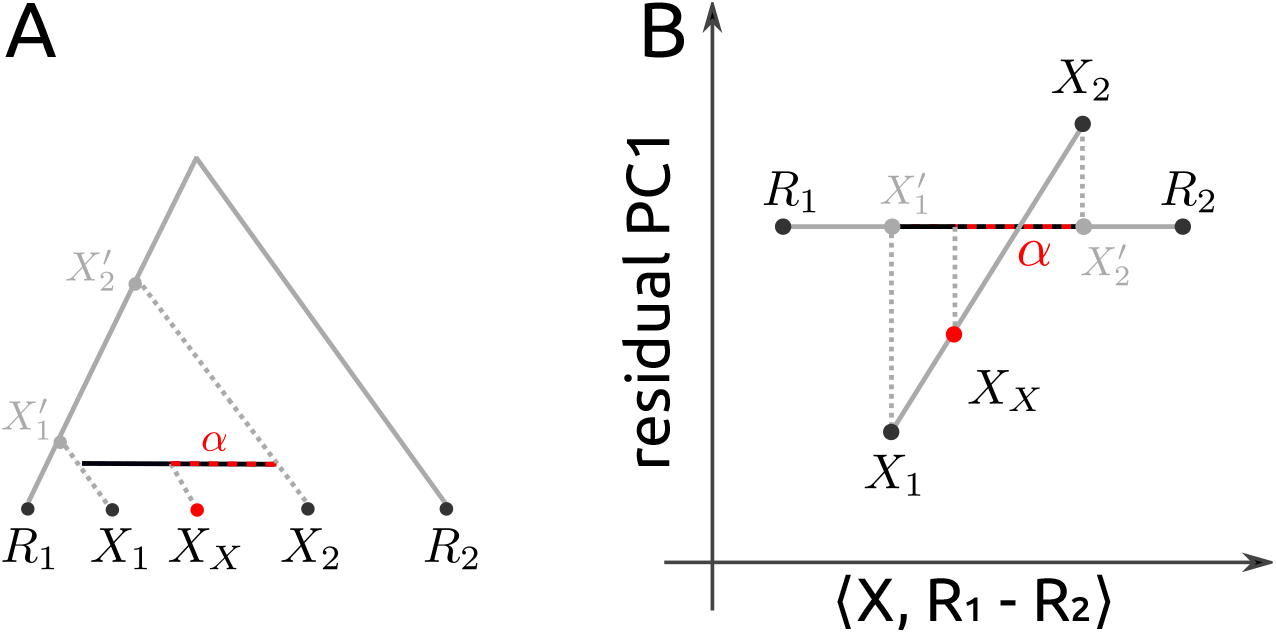
Admixture proportion estimates. A: Visualization of the admixture graph scenario used to estimate the ancestry proportion *α* contributed from *X*_1_ to *X_X_*, using references *R*_1_ and *R*_2_. The full grey line corresponds to the projection axis, and the dotted grey lines corresponds to the branches ignored in the projections. The admixture proportion *α* corresponds to the length of the dashed red line relative to the black line between *X*_1_ and *X*_2_. B: The same scenario, but in Euclidean space, *X*_1_, *X*_2_ and *X_X_* align on a line both in the (low-dimensional approximation of the) residual space and on the projection axis.

The PCA-like projection can be used to visualize admixture proportions, as the horizontal position of *X_X_* relative to *X*_1_ and *X*_2_ (red dashed line vs black line) directly represents the estimated admixture proportion *α*. In addition, the residuals can be used to verify assumptions of the admixture graph model. In particular, since *X_X_* arises as a linear combination of *X*_1_ and *X*_2_, if admixture is recent we might expect the three populations to be collinear; if they are not this means that either of the populations experienced gene flow from some other population which might bias results (Petr et al., 2019).

In addition, the external tree branches 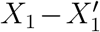 and 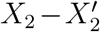 are disjoint which means they should be orthogonal. One a one-dimensional residual plot (Figure 6B) this can not be verified, but the statistic

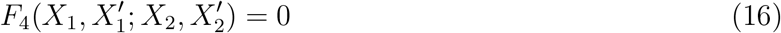

can be calculated for all samples.

#### 4.5.1 Example

I use the World data set as an example, using Sardinian and Mbuti as references populations (Figure 5B). The data is the same as in the PCA (Figure 5A), but it is now rotated such that the line between the reference population (black line in Figure 5A) now corresponds to the x-axis. For any pair of populations *X*_1_ and *X*_2_, their horizontal distance reflects *F*_4_(Sardinian, Mbuti; *X*_1_, *X*_2_) and the relative horizontal distance corresponds exactly to *F*_4_-ratio admixture estimates. For many sets of populations, this is of course not sensible, and just looking at the first PC of the residual shows many examples where the populations are not collinear. For example, if we just focus on the x-axis, the South American Surui appear between Papuans and Georgians. But since the Surui clearly are not on the line between Papuans and Georgians, this plot reveals that this should not be interpreted as the result of admixture.

## 5 Discussion

Particularly for the analysis of human genetic variation with a large number of individuals with heterogeneous relationships, *F*-statistics are a powerful tool to describe population genetic diversity. Here, I show that the geometry of *F*-statistics (Oteo-Garcia and Oteo, 2021) leads to a number of simple interpretations of *F*-statistics on a PCA-plot.

### 5.1 The Geometry of Admixture

Previous interpretation of PCA in the context of population genetic models have focused on explicit models and aimed at directly interpreting the PCs in terms of population genetic parameters (Cavalli-Sforza and Piazza, 1975; Novembre and Stephens, 2008; François et al., 2010; François and Gain, 2021). My interpretation here is different in that the utility of PCA is to simplify the geometry of the data. One consequence is that the results here are not directly impacted by sample ascertainment, sample sizes or the number of principal components analyzed, which are common concerns in the interpretation of PCA. However, a very skewed sampling distribution will increases the likelihood that more, or different PCs will have to be included in the analysis.

The two data sets I analyzed here suggest that two PCs (for the World data set) respectively three PCs (for the Western Eurasian data) already provide a very good approximation for *F*_4_-statistics (Figures 3E, 5E), reflecting the observation that frequently the first few PCs provide a good approximation of population structure. On the other hand, for admixture *F*_3_-statistics, more PCs are needed (Figure 3BC), likely because these statistics do contain a term measuring local variation in the putative admixed population (Peter, 2016), which is typically often incorporated on higher PCs. Thus, I would also expect individual-based analyses and *F*_4_-statistics between closely related populations to require substantially more PCs.

My focus on the geometry of the data allows for direct and quantitative comparisons between *F*-statistic-based results and PCA biplots. As PCA is often ran in an early step in data analysis, this may aid in generation of hypotheses that can be more directly evaluated using generative models, typically using a lower number of populations. It also allows reconciling apparent contradictions between *F*-statistics and PCA-plots. In many cases, differences between the two data summaries will be due to variation on higher PCs. In this case, plotting additional PCs, or further subsetting the data to a more local set of populations seems prudent.

### 5.2 Assumptions

The other cause for disagreements between *F*-statistics and PCA are differences in assumptions. The version of PCA I used for my analyses was chosen such that the similarities to *F*-statistics are maximized. In particular, I assume here that i) we have no missing data, ii) SNPs are equally weighted and iii) that individuals can be grouped into populations and iv) we use estimated allele frequencies. In contrast, most data analyses have to grapple with missing data, SNP are often weighted according to their allele frequency and observed, individual-level genotypes are used as the basis of PCA.

#### 5.2.1 Missing data

The liability of PCA to missing data is a well-studied problem and a number of algorithms for imputing missing data have been proposed (e.g. Hastie et al., 2015; Meisner and Albrechtsen, 2018; Meisner et al., 2021). Calculating *F*-statistics first is yet another way to impute data for PCA that works well if missingness is low (Meisner et al., 2021). In contrast, missing data in *F*-statistics is handled by estimating a standard error using resampling across the genome (Patterson et al., 2012), which does not distinguish between biological and sampling variation. These strategies are distinct, but not unique to the relative approaches and a PCA-like decomposition from *F*-statistics is commonly applied using MDS (e.g. Fu et al., 2016). The theory developed here suggests that developing a PCA-based *F*-statistics framework might be more powerful, as related populations could be used to fill in missing data.

#### 5.2.2 Normalization

In PCA, SNPs are typically normalized to have expected variance of one; a step that is omitted in calculating *F*-statistics (Patterson et al., 2006). The *F*-statistic framework assumes that each SNP is an identically-distributed (but not independent) random variable. which holds regardless of weighting. Thus, normalization of SNPs is largely a matter of convention, for *F*-statistics the dependency on additional samples (through mean allele frequencies) is often unwanted, but could be advantageous for tools that aim to do joint inference from many *F*-statistics such as `qpAdm` (Patterson et al., 2012; Harney et al., 2021). As genetic differentiation between human populations is low, the normalization used may matter little in practice, but should be explored in future work (Felsenstein, 1973).

#### 5.2.3 Estimated vs. observed allele frequencies

The third difference between *F*-statistics and standard PCA is on the usage of estimated allele frequencies versus individual-based genotypes. The fact that PCA does not distinguish between sample-based errors and the underlying structure is a well-known drawback of standard PCA, and applying the theory presented here to individual-based PCA would result in *F*-statistics that incorporate some sampling noise. Probabilistic PCA is one class of approaches that aim to separate the population structure from sampling noise (e.g. Agrawal et al., 2020). It seems likely that probabilistic PCA would yield a representation of the data that corresponds more closely aligned with *F*-statistics than regular PCA.

#### 5.2.4 Individual vs. population-based analyses

The final issue is that PCA is commonly run on individual-based data, whereas *F*-statistics often group individuals into populations. This is not necessary; population-based PCA has been the default in the past (Cavalli-Sforza et al., 1994), and *F*-statistics are often applied to individuals (e.g. Green et al., 2010; Massilani et al., 2020; Yang et al., 2020). Thus, more commonly the choice between individual-based and population-based approaches is in the goal of an analysis. Grouping individuals into populations introduces a major assumption, that can e.g. be justified using an individual-based PCA. In particular, since *F*-statistics assume individuals are randomly drawn from a population, they should form tight clusters on an individual-based PCA-plot, otherwise population substructure becomes a possible alternative model for negative *F*_3_-statistics and non-zero *F*_4_-statistics (Peter, 2016).

#### 5.2.5 Summary

The version of PCA used here differs from that proposed by Patterson et al., 2006, and thus some care will be required to directly extend the interpretations developed here to individual-based PCAs. However, the differences are largely due to conventions, and particularly for studies where the description of population structure is a major focus, results might be easier to interpret if conventions regarding missing data, normalization and estimation of allele frequencies are used consistently between *F*-statistics and PCA.

### 5.3 The Apportionment of Human Diversity

Most genetic variation in humans is shared between all of us, but the around 15% that can be explained by population structure can be leveraged to study our history and diversity in great detail (Lewontin, 1972; Cavalli-Sforza et al., 1964; Reich, 2018). For some data sets it is possible to predict an individuals’ origin at a resolution of a few hundred kilometers (Novembre et al., 2008; Leslie et al., 2015), and direct-to-consumer-genetics companies are using this variation to analyze the genetic data of millions of customers.

However, understanding, conceptualizing and modeling this variation is far from trivial, and misconstrued models of human genetic differentiation have repeatedly been used to justify racist, eugenic and genocidal policies. Lewontin’s landmark 1972 paper on the apportionment of human genetic diversity was one of the first to quantify how little of between-population genetic variation could be attributed to “racial” continental-scale groupings. Over the last five decades, this view has been corroborated, refined and extended many times (Cann et al., 1987; Cavalli-Sforza et al., 1994; Barbujani et al., 1997; Rosenberg et al., 2002).

From a practical perspective, formulating hypotheses and designing studies in terms of discrete populations with “uniform” genetic backgrounds is often sensible, as it enables e.g. prediction of phenotypes (Berg et al., 2019; Yair and Coop, 2021), inference of demographic parameters and schematic models of human genetic history (Patterson et al., 2012). This is also the case for *F*-statistics, which are motivated by trees. They assume that populations are discrete, related as a graph, and that gene flow between populations is rare (Patterson et al., 2012; Harney et al., 2021). However, these simplification do come at a cost, both in terms of model violations that may invalidate statistical results, and in terms of deemphasizing people that do not rigidly fall into predefined genetic groups.

In many parts of the world, and particularly at more local scales, distinctions between populations begin to blur, and everyone could be considered admixed to some degree (Pickrell and Reich, 2014). This provides a challenge for interpretation; as most *F*_3_ and *F*_4_-statistics will indicate departures from treeness. A naive interpretation of the *F*-statistics from my Eurasian example (Figure 3A) would identify a substantial fraction of Europeans as (significantly) admixed between Finnish and Sardinians. In contrast, PCA reveals that the variation in this data set is not due to a single event, and so an arguably better description of the data set is one where Finnish and Sardinians lie on opposite ends of a more gradually structured population.

This continuum between admixture and population structure becomes more problematic when interpreting tools that use multiple *F*-statistics to build compound models, such as `qpGraph` (Lazaridis et al., 2014) and `qpAdm` (Harney et al., 2021). One issue with these approaches is that they are usually restricted to at most a few dozen populations. As ancient DNA data sets now commonly include thousands of individuals, analysts are faced with the challenge of which data to include. A common approach is to sample a large number of distinct models, and retain the ones that are compatible with the data. However, as both `qpGraph` and `qpAdm` assume that gene flow is rare and discrete, selecting sets of populations that did experience little gene flow will provide good fits. The PCA-based interpretation of *F*-statistics offers an alternative that trades interpretability for robustness, as I do not need to assume that gene flow is rare.

## 6 Supplementary File Description

Both supplementary files give the individual-identifier, sex and population label for each sample.

## A Derivations

Depending on a readers’ background in linear algebra, these results may appear elementary; I include them here for reference and because they were not obvious to me at the onset of this project.

### *F*-statistics are invariant under a change-of-basis

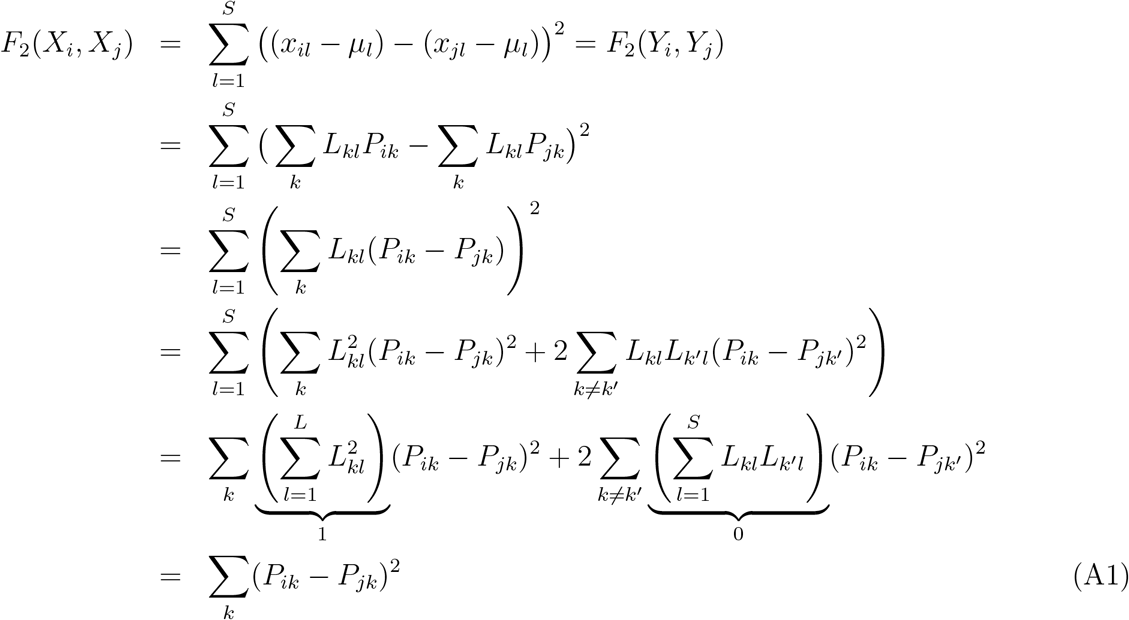

In summary, the first row shows that *F*_2_ on the centered data will give the same results (as distances are invariant to translations), in the second row we apply the PC-decomposition. The third row is obtained from factoring out *L_lk_*. Row four is obtained by multiplying out the sum inside the square term for a particular *l*. We have *k* terms when for 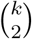 terms for different *k*’s. Row five is obtained by expanding the outer sum and grouping terms by *k*. The final line is obtained by recognizing that **L** is an orthonormal basis; where dot products of different vectors have lengths zero.

Note that if we estimate *F*_2_, unbiased estimators are obtained by subtracting the population-heterozygosities *H_i_, H_j_* from the statistic. As these are scalars, they do not change above calculation.

### The region of negative *F*_3_-statistics is a *n*-ball

Without loss of generality, assume that *X*_1_ = (*r*, 0, 0,…) and *X*_2_ = (−*r*, 0, 0,…), and let us assume that *X_x_* has coordinates (*x*_1_, *x*_2_,…, *x_S_*) Assuming *F*_3_(*X_x_*; *X*_1_, *X*_2_) = 0, equation 13 becomes

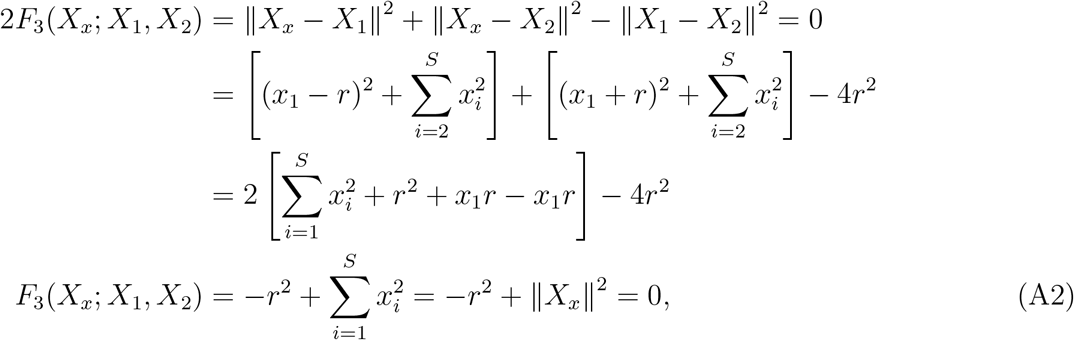

which is the equation of a *n*-sphere with radius *r* and center at the origin, as assumed from the placing of *X*_1_ and *X*_2_. Now, assume that *F*_3_ is negative, i.e. *F*_3_(*X_x_*; *X*_1_, *X*_2_) = −*k* < 0. Moving *r*^2^ to the left we obtain

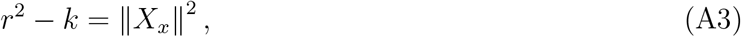

which is another *n*-sphere with a smaller radius, showing that all points inside the *n*-sphere will have negative *F*_3_-values.

### If a population lies outside the circle of this *n*-Sphere in any 2D-projection *F*_3_ is positive

Assume the center of the *n*-sphere 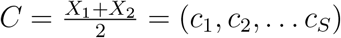, and *X_x_* = (*x_i_*,… *x_S_*). Then,

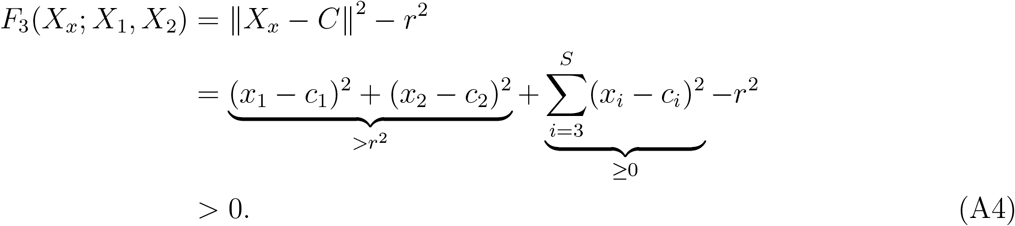

The condition (*x*_1_ – *c*_1_)^2^ + (*x*_2_ – *c*_2_)^2^ > *r*^2^ is satisfied whenever *X_x_* is outside the circle obtained from projecting the *n*-sphere on the first two dimensions. An analogous argument applies for any low-dimensional representation.

